# Metal complexes and conjugation: Harnessing the power of cobalt complexes to curtail plasmid transfer

**DOI:** 10.1101/2023.11.24.568573

**Authors:** Ilyas Alav, Parisa Pordelkhaki, Pedro Ernesto de Resende, Hannah Partington, Simon Gibbons, Rianne Lord, Michelle M.C. Buckner

## Abstract

**Background:** Antimicrobial resistance genes (ARG), such as extended spectrum β-lactamase (ESBL) and carbapenemase genes, are commonly carried on plasmids. Plasmids can transmit between bacteria, disseminate globally, and cause clinically important resistance. Therefore, targeting plasmids could reduce ARG prevalence, and restore the efficacy of existing antibiotics. Here, we assessed the effect of four previously characterised bis(*N*-picolinamido)cobalt(II) complexes on the conjugative transfer of plasmids in *Escherichia coli* and *Klebsiella pneumoniae*.

**Methods:** Liquid broth and solid agar conjugation assays were used to measure complex activity on four plasmids in *E. coli*. Additionally, the effect of cobalt complexes was tested on the transmission of the fluorescently tagged extended spectrum β-lactamase encoding pCT*gfp* plasmid in *E. coli* and carbapenemase encoding pKpQIL*gfp* plasmid in *K. pneumoniae*, using flow cytometry.

**Results:** Antimicrobial susceptibility testing of cobalt complexes revealed no antibacterial activity. The cobalt complexes significantly reduced conjugative transfer of RP4, R6K, and R388 plasmids on solid agar in *E. coli* and pKpQIL*gfp* transmission in *K. pneumoniae.* None affected conjugative transfer of pKM101 or transmission of fluorescently tagged pCT in *E. coli*. The cobalt complexes had no effect on plasmid persistence, suggesting that they target conjugation rather than plasmid prevalence.

**Conclusions:** To the best of our knowledge, this is the first study to report reduced transmission of clinically relevant plasmids with cobalt complexes. These cobalt complexes are not cytotoxic towards mammalian cells and are not antibacterial, therefore they could be optimised and employed as conjugation inhibitors to reduce prevalence of AMR and/or virulence genes in animals and humans.

**Significance:** Antimicrobial resistance is a growing problem that poses a significant threat to modern medicine. Some of the most problematic resistance genes are carried on genetic elements, called plasmids, that can spread between bacteria. While our understanding of the mechanisms and drivers of gene transfer amongst bacteria is increasing, we lack effective tools to slow down/control these processes. Here we demonstrate for the first time that novel cobalt-based compounds have anti-plasmid activity on a subset of *E. coli* plasmids, and are extremely potent in *K. pneumoniae* carrying a clinical carbapenem-resistance plasmid, without impacting plasmid maintenance. This finding forms the foundations of a potential strategy to control the transfer of genes within Gram-negative bacteria, which has implications for AMR and virulence.

## Introduction

Antimicrobial resistance (AMR) is a global public health crisis that jeopardises our ability to treat infectious diseases (1, 2). The threat of AMR has been further compounded by the dissemination of antimicrobial resistance genes (ARGs) by mobile genetic elements such as plasmids, which can carry multiple genes encoding proteins that confer resistance to a wide range of clinically relevant antibiotics (3, 4). Worryingly, opportunistic pathogens like multidrug-resistant (MDR) *Enterobacteriaceae*, such as *Escherichia coli* and *Klebsiella pneumoniae*, often harbour plasmids that carry ARGs coding for extended-spectrum β-lactamases (ESBLs e.g., CTX-M) and carbapenemases (e.g., KPC), which confer resistance to β-lactam and carbapenem antibiotics, respectively (5–8). Hence, infections caused by MDR *Enterobacteriaceae* have become increasingly difficult to treat due to dwindling treatment options (9). Consequently, patients with carbapenem-resistant *Enterobacteriaceae* infections face a significantly greater risk of death compared to patients with carbapenem susceptible *Enterobacteriaceae* infections (10). In 2017, the World Health Organisation (WHO) recognised the threat posed by carbapenem-resistant and ESBL-producing *Enterobacteriaceae* and designated this pathogen group as a critical priority for which novel drugs are needed (11). Therefore, there is an urgent need to develop new strategies to treat these infections, including making carbapenem-resistant and ESBL-producing *Enterobacteriaceae* susceptible to existing drugs and preventing the spread of AMR between bacteria.

ARGs are commonly found in conjugative AMR plasmids that can be readily shared between bacteria that occupy the same environmental niche (12–14). The transmission of AMR plasmids between different species of *Enterobacteriaceae* has been well documented in both clinical and environmental settings (15–17). Once *Enterobacteriaceae* acquire ESBL- or carbapenemase-producing AMR plasmids, they can become MDR and extremely challenging to eradicate when they cause infections (18). There are several different factors that contribute to the prevalence and spread of AMR plasmids, including high conjugation rates, increased plasmid copy number, reduced fitness cost of plasmid carriage due to compensatory mutations, and successful plasmid and clonal group interplay (19–24). Owing to the potential to transfer multiple ARGs simultaneously, their high mobility and persistence, and the significant impact they have on treatment options, AMR plasmids are a serious threat to both animal and human health.

A potential strategy to overcome the threat of carbapenem-resistant and ESBL-producing *Enterobacteriaceae* is by reducing the prevalence of AMR plasmids (25). This could restore the efficacy of existing well-tolerated drugs and reduce the necessity of using of more toxic alternatives. Compounds that target plasmids can work by removing the plasmid from a population by reducing plasmid stability (plasmid curing) and/or interfering with the conjugation process to prevent the transfer of a plasmid to a new host (25–27). Previous studies have identified compounds with plasmid curing/conjugation inhibiting activity, ranging from phytochemicals to clinically approved drugs (28–31). Except for clinically approved drugs, the majority of the previously described compounds such as biocides, DNA intercalating agents, and detergents are either toxic or have not been tested in clinically relevant AMR plasmids of *Enterobacteriaceae* (25).

Cobalt (Co) is a trace element in the body and is essential for many biological processes, and whilst excess amounts or deficiency of the metal can induce undesired effects (32). Cobalt complexes have found usage as anticancer, antiviral, and antimicrobial agents (33, 34). In particular, Co(II) and Co(III) complexes have been reported to have high activities against *Staphylococcus aureus* and *E. coli* (35), and fungal strains (36). Previously, tris(*N*-picolinamido)cobalt(III) complexes were reported to have antibacterial activity against *Pseudomonas* and *E. coli* (35). However, Ghandhi *et al* (37) reported the ESKAPE screening (CO-ADD; Community for Open Antimicrobial Drug Discovery, The University of Queensland, Australia) of a range of bis(*N*-picolinamido)cobalt(II) complexes and confirmed they had minimal antibacterial activity and importantly, no activity against mammalian cell lines (37). These results suggest the oxidation state of the cobalt complexes could induce differences in their bacterial mode of action. Here, we evaluated the activity of some of these cobalt complexes lacking antibacterial activity for their ability to reduce conjugation of various plasmids in *E. coli*. They were found to inhibit the transmission of the carbapenemase drug resistance plasmid pKpQIL in *K. pneumoniae*. This study demonstrates the viability of using non-toxic cobalt complexes as a strategy to reduce plasmid conjugation.

## Materials and method

### Strains and plasmids

All bacterial strains and plasmids used in this study are listed in Tables 1 and 2, respectively. Unless stated otherwise, all strains were grown in Luria-Bertani (LB) broth/broth with agar (Merck, Germany) at 37 °C with aeration (liquid cultures).

**Table 1.**
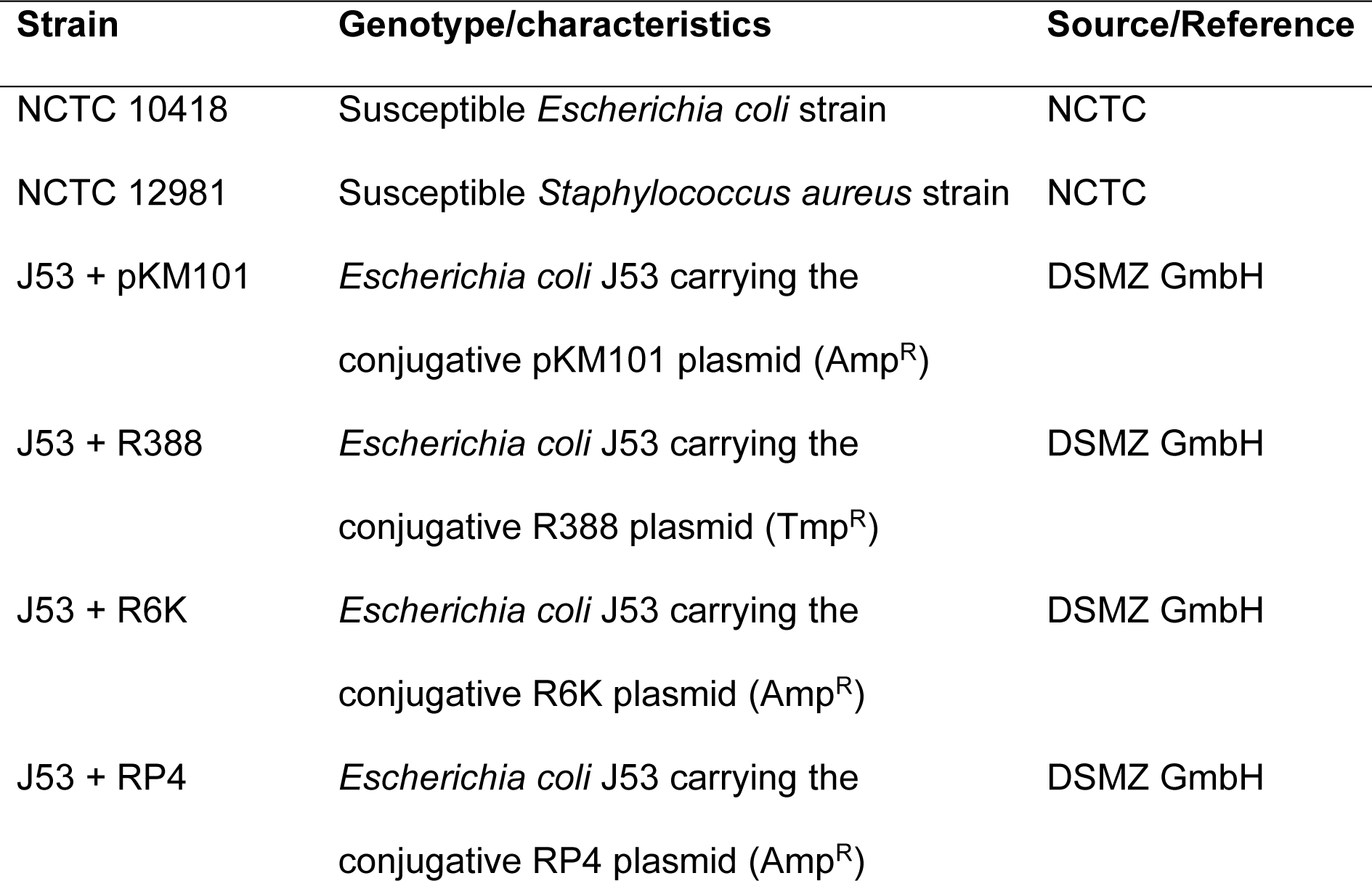
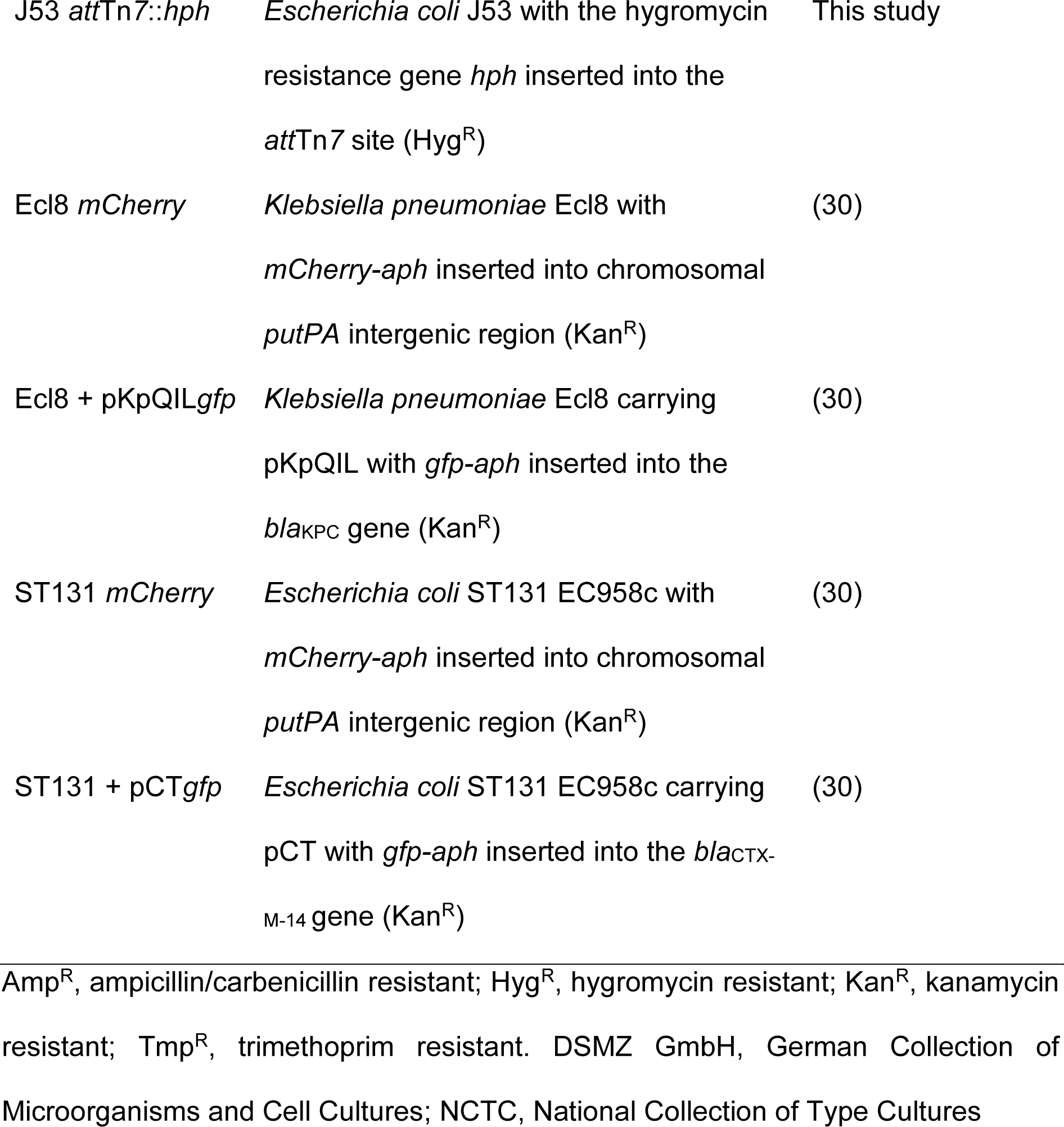
List of bacterial strains used in this study.

**Table 2.** List of plasmids used in this study.

| Plasmid | Description | Source/Reference |
| --- | --- | --- |
| pKM101 | IncN plasmid with MOB<sub>F11</sub> type relaxase and ampicillin resistance gene | DSMZ GmbH |
| R388 | IncW plasmid with MOB<sub>F11</sub> type relaxase and trimethoprim resistance gene | DSMZ GmbH |
| R6K | IncX2 plasmid with MOB<sub>P3</sub> type relaxase and ampicillin streptomycin and resistance genes | DSMZ GmbH |
| RP4 | IncP plasmid with MOB<sub>P11</sub> type relaxase and ampicillin resistance gene | DSMZ GmbH |
| pSIM18 | Plasmid used as a template to amplify the hygromycin resistance gene | (38) |
| pSLTS | Plasmid encoding arabinose inducible lambda red recombination system to facilitate homologous recombination | (39) |
| pCT*gfp* | IncK plasmid pCT with *gfp-aph* inserted into the *bla*<sub>CTX-M-14</sub> gene | (30) |
| pKpQIL*gfp* | IncFII<sub>K2</sub> plasmid pKpQIL with *gfp-aph* inserted into the *bla*<sub>KPC</sub> gene | (30) |

DSMZ GmbH; German Collection of Microorganisms and Cell Cultures

### Determination of antibacterial susceptibility

The broth microdilution method was used to determine the minimum inhibitory concentrations (MICs) of the cobalt complexes according to Clinical and Laboratory Standards Institute guidance (40). *E. coli* NCTC 10418 and *S. aureus* NCTC 12981 were used as quality control strains (Table 1). The MIC values were recorded as the lowest concentration at which no bacterial growth was detected. All MICs were carried out using three biological replicates.

### Growth kinetic assays

The impact of the cobalt complexes on bacterial growth was determined as previously described (41). Briefly, overnight cultures (∼10^9^ CFU/mL) of test strains were diluted to a starting inoculum of 10^6^ CFU/mL in a 96-well flat bottom plate (Corning, USA). Where appropriate, the test strains were diluted in LB broth supplemented with the cobalt complexes or DMSO vehicle control to a final concentration of 100 µg/mL. Growth was monitored at OD_600_ at 30 min intervals for 12 h using the FLUOstar OMEGA plate reader (BMG Labtech, Germany). Data shown are the mean OD_600_ per time point ± standard deviation of three independent experiments, each with three biological replicates.

### Construction of the hygromycin resistant *Escherichia coli* J53 recipient strain

To obtain a hygromycin resistant *E. coli* J53 strain to be used as a recipient for conjugation assays, the *hph* gene encoding hygromycin B phosphotransferase was inserted into the phenotypically neutral *att*Tn*7* site (42) using the arabinose inducible recombineering plasmid pSLTS as described previously (39). Firstly, the *hph* gene was amplified from the pSIM18 plasmid using primers that have flanking 40 bp homology to the *att*Tn*7* site in *E. coli* (Table S1). The arabinose inducible recombineering plasmid pSLTS was electroporated into *E. coli* J53 with subsequent electroporation of the PCR-amplified hygromycin resistance cassette. Successful recombinants were selected on LB agar supplemented with 150 µg/mL hygromycin. PCR and Sanger sequencing (Eurofins Genomics, UK) using primers that bind upstream and downstream of the recombination site (Table S1) were used to verify successful insertion of the *hph* gene at the desired genomic locus. Antimicrobial susceptibility testing was also used to verify the hygromycin resistant phenotype (MIC >512 µg/mL).

### Liquid broth conjugation assay

The donor *E. coli* J53 strain with R388, pKM101, RP4 or R6K was paired with the hygromycin resistant recipient strain *E. coli* J53 *att*Tn*7*::*hph*. The liquid broth conjugation assays were performed as previously described with minor modifications Donor and recipient cultures were grown overnight, sub-cultures were prepared in 5 mL LB broth (1% inoculum) and grown to an OD_600_ of ∼0.5. A 1 mL volume of culture was pelleted, and media were replaced with LB broth to normalise the OD_600_ to 0.5. Equal volumes of donor and recipient strains were mixed to give a donor-to-recipient ratio of 1:1. Cultures were diluted 1:5 in LB broth containing a final concentration of 100 µg/mL of cobalt complexes or 100 µg/mL DMSO as vehicle control and these were incubated statically at 37 °C for 4 h. Corresponding dilutions were plated on selective media and incubated at 37 °C overnight. Transconjugant colonies carrying RP4, R6K or pKM101 were selected on LB agar supplemented with 150 µg/mL hygromycin B (PhytoTech Labs, USA) and 100 µg/mL carbenicillin (Merck, Germany). Transconjugant colonies carrying R388 were selected on LB agar supplemented with 150 µg/mL hygromycin B and 10 µg/mL trimethoprim (Merck, Germany). Conjugation frequencies (CF) were calculated using the following formula:

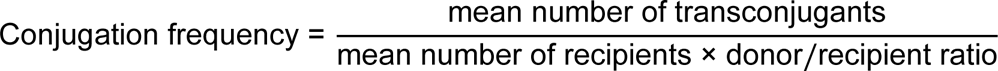

Data shown are the mean ± standard deviation of three independent experiments, each carried out with four biological replicates.

### Solid agar conjugation assay

The donor *E. coli* J53 strains with R388, pKM101, RP4 or R6K were paired with the hygromycin resistant recipient strain *E. coli* J53 *att*Tn*7*::*hph*. The solid agar conjugation assay was performed as previously described with minor modifications Briefly, a 1 mL volume of overnight cultures of donor and recipient cells was pelleted, washed with LB broth, and the OD_600_ was adjusted to 0.5. Equal volumes of donor and recipient cells were mixed to give a donor-to-recipient ratio of 1:1. Then, 5 µL of this mixture was placed on top of 96-well round bottom plates (Corning, USA) containing 150 µL LB agar supplemented with 100 µg/mL of cobalt complexes or 100 µg/mL DMSO as vehicle control. Conjugation was carried out for 4 h at 37 °C without agitation. Bacteria were resuspended in 150 µL phosphate-buffered saline (PBS), and corresponding dilutions were plated on selective media as described above and incubated at 37 °C overnight. Conjugation frequencies were calculated as for the liquid conjugation assay. Data shown are the mean ± standard deviation of three independent experiments, each carried out with four biological replicates.

### Measurement of plasmid transmission by flow cytometry

The transmission of pCT*gfp* in *E. coli* ST131 EC958c and pKpQIL*gfp* in *K. pneumoniae* ECl8 was measured by flow cytometry as previously described with minor modifications (30). Briefly, 1 mL overnight cultures of donor (*E. coli* with pCT*gfp* or *K. pneumoniae* with pKpQIL*gfp*) and recipient (*E. coli* or *K. pneumoniae* with chromosomal mCherry) strains were pelleted, washed in PBS, and diluted to an OD_600_ of 0.5. Equal volumes of donor and recipient strains were mixed to give a donor-to-recipient ratio of 1:1. A 20 µL volume of the donor-recipient mix was inoculated into 180 µL of LB broth supplemented with a final concentration of 100 µg/mL of cobalt complexes or 100 µg/mL DMSO as vehicle control in a 96-well round bottom plate (Corning, USA). The plate was incubated at 37 °C with gentle agitation (∼100 rpm) for 4 h. Following incubation, 20 µL was removed and serially diluted 1:1000 in filter-sterilised Dulbecco’s PBS (Merck, Germany). Samples were analysed on the Attune NxT acoustic focusing flow cytometer with Autosampler (Thermo Scientific, USA). GFP emission was collected using the BL1-H channel and the mCherry emission was collected using the YL2-H channel. Plasmid transmission was measured by quantifying the number of green fluorescent protein (GFP)-positive bacteria (donor), mCherry-positive bacteria (recipient), and GFP-positive/mCherry-positive bacteria (transconjugants). Gating strategies were exactly as previously described (30). Plasmid transmission was calculated as the number of dual fluorescent bacterial events divided by the total bacterial events relative to the DMSO control. Data shown are the mean ± standard deviation of three independent experiments, each carried out with four biological replicates.

### Plasmid persistence

Overnight cultures of *E. coli* J53 carrying RP4, R6K, R388, or pKM101, *E. coli* ST131 EC958c carrying pCT*gfp*, and *K. pneumoniae* Ecl8 carrying pKpQIL*gfp* were grown. Sub-cultures were prepared in 5 mL LB broth (5% inoculum) and grown to an OD_600_ of ∼0.6. A 1 mL volume of culture was pelleted, and media were replaced with LB broth to normalise the OD_600_ to 0.5. A 5 µL volume of culture was inoculated into 195 µL of LB broth supplemented with a final concentration of 100 µg/mL cobalt complexes or 100 µg/mL DMSO as vehicle control in a 96-well round bottom plate (Corning, USA). The plate was then incubated at 37 °C for 24 h without agitation. Following 24 h incubation, each well was serially diluted to 10^-6^ in PBS. A 10 µL volume of the diluted culture was then used to passage the cells in 190 µL LB broth supplemented with a final concentration of 100 µg/mL cobalt complexes or 100 µg/mL DMSO as vehicle control in a 96-well round bottom plate for a further 24 h incubation. Corresponding dilutions were plated 24 h and 48 h post incubation on both selective media and non-selective media and incubated at 37 °C overnight. The plasmids RP4, R6K, and R388 were selected on 100 µg/mL carbenicillin, R388 on 10 µg/mL trimethoprim, and pCT*gfp* and pKpQIL*gfp* on 50 µg/mL kanamycin. Percentage of plasmid persistence was calculated as

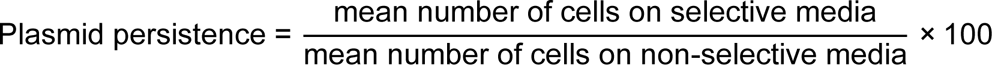

### Statistical analysis

Unpaired *t*-tests were used for statistical analysis with GraphPad Prism version 10 for MacOS, San Diego, California USA, http://www.graphpad.com. Only P-values less than or equal to 0.05 were considered statistically significant.

## Results

### Cobalt complexes are not antibacterial

Firstly, the susceptibility of the test strains (Table 1) to the cobalt complexes Co4, Co5, Co6, and Co8 (Fig. S1) were determined to detect any antibacterial activity and to identify a suitable concentration to evaluate their effect on plasmid conjugation. In agreement with the previous study (37), none of the tested cobalt complexes exhibited antibacterial activity against the strains tested up to a concentration of 512 µg/mL (Table 3). Based on the minimum inhibitory concentration results, the impact of the cobalt complexes on plasmid transfer was tested at 100 µg/mL. Bacterial growth kinetics in the presence of 100 µg/mL cobalt complexes compared to 100 µg/mL DMSO vehicle control showed that none had any significant adverse effects on bacterial growth over 12 h (Fig. S2).

**Table 3.** Susceptibility of the test strains to the bis(*N*-picolinamido)cobalt(II) complexes.

| Strain | MIC (µg/mL) |  |  |  |
| --- | --- | --- | --- | --- |
|  | Co4 | Co5 | Co6 | Co8 |
| *S. aureus* NCTC 12981 | >512 | >512 | >512 | >512 |
| *E. coli* NCTC 10418 | >512 | >512 | >512 | >512 |
| *E. coli* J53 *attTn7::hph* | >512 | >512 | >512 | >512 |
| *E. coli* J53 with RP4 | >512 | >512 | >512 | >512 |
| *E. coli* J53 with R6K | >512 | >512 | >512 | >512 |
| *E. coli* J53 with R388 | >512 | >512 | >512 | >512 |
| *E. coli* J53 with pKM101 | >512 | >512 | >512 | >512 |
| *K. pneumoniae* Ecl8 *mCherry* | >512 | >512 | >512 | >512 |
| *K. pneumoniae* Ecl8 with pKpQILgfp | >512 | >512 | >512 | >512 |
| *E. coli* ST131 EC958c *mCherry* | >512 | >512 | >512 | >512 |
| *E. coli* ST131 EC958c with pCTgfp | >512 | >512 | >512 | >512 |

Minimum inhibitory concentration (MIC) values shown are the median of three biological replicates.

### Cobalt complexes affect conjugative plasmid transfer differently in liquid and solid mating

The effect of the cobalt complexes on plasmid conjugation was tested using plasmids belonging to different incompatibility groups to investigate the specificity of conjugation inhibition in *E. coli* (Table 2). Plasmid conjugation frequencies (CF) are also known to differ in liquid broth and on solid surfaces (45). Therefore, the effect of cobalt complexes on conjugative plasmid transfer was tested using both liquid broth and agar mating experiments. In agreement with previous studies (45, 46), the IncP plasmid RP4, the IncX2 plasmid R6K, the IncW plasmid R388 and the IncN plasmid pKM101 displayed higher CFs on solid agar compared to liquid broth mating (Fig. 1 and Table S2).

**Figure 1.**
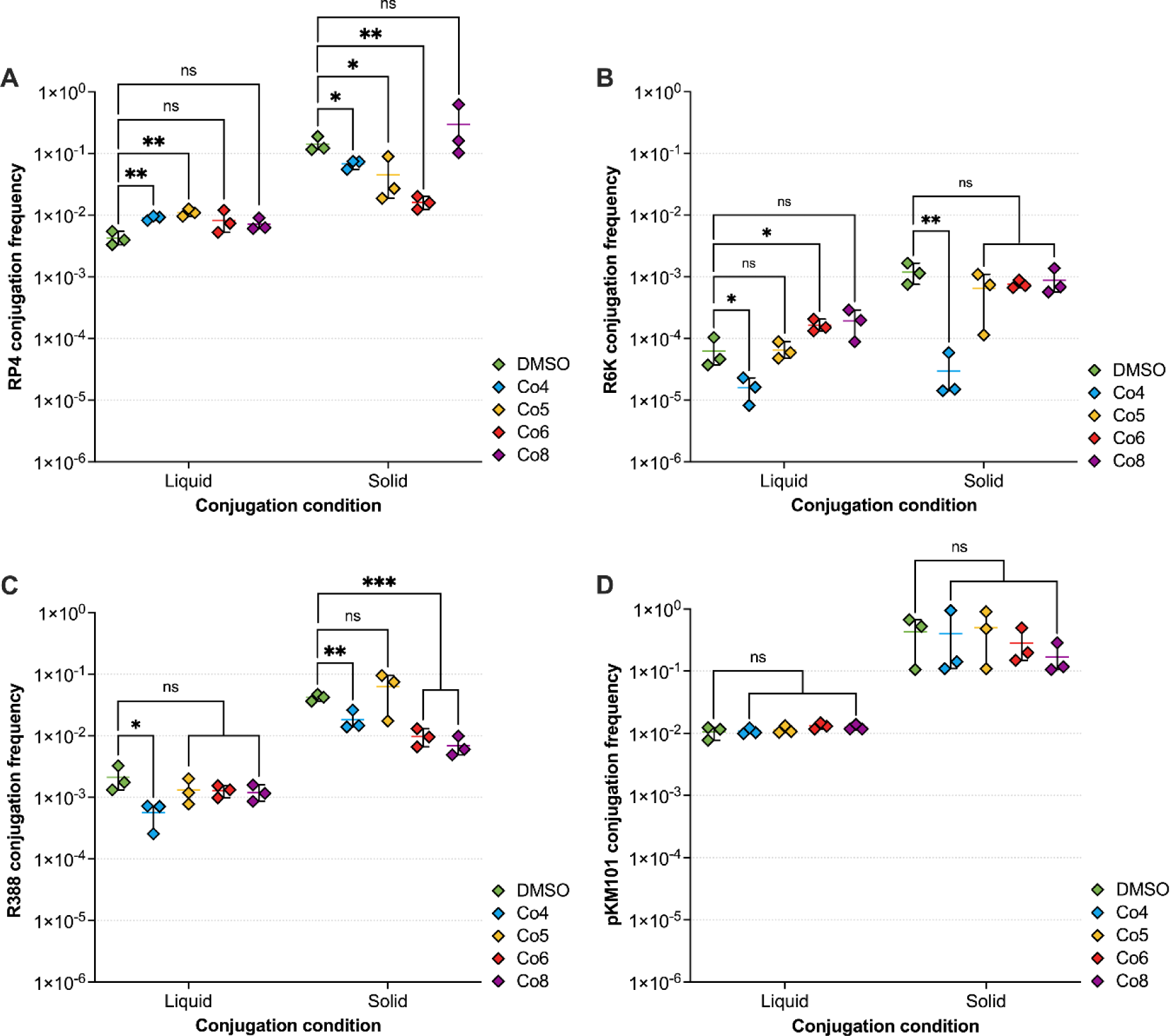
The effect of cobalt complexes on the conjugation frequencies of plasmids with different incompatibility groups in liquid LB broth and on LB agar. Conjugation frequencies of **(A)** the IncP plasmid RP4, **(B)** the IncX2 plasmid R6K, **(C)** the IncW R388, and **(D)** the IncN plasmid pKM101, from *E. coli* J53 to hygromycin resistant *E. coli* J53 *att*Tn*7*::*hph* in the presence of 100 µg/mL DMSO vehicle control or 100 µg/mL cobalt compound after four-hour incubation. Data shown are the mean ± standard deviation of three independent experiments, each carried out with four biological replicates. Cobalt complexes that significantly affected conjugation frequency compared to DMSO control are indicated with * (*p* ≤ 0.05), ** (*p* ≤ 0.01) or *** (*p* ≤ 0.001). ns, not significant.

Interestingly, the cobalt complexes affected plasmid CFs differently depending on the conjugation condition. Co4 and Co5 significantly increased the CF of RP4 in liquid broth mating, whilst in solid agar mating they both significantly reduced CF of RP4 (Fig. 1A) from 1.42×10^-1^ in DMSO control to 6.79×10^-2^ in Co4 (*p* = 0.0186) and 4.49×10^-2^ in Co5 (*p* = 0.0197) (Table S2). Whilst Co6 had no impact on the CF of RP4 in liquid broth (Fig. 1A), it significantly reduced CF in plate mating from 1.42×10^-1^ in DMSO control to 1.61×10^-2^ in Co6 (*p* = 0.0029) (Table S2). Co8 had no effect on RP4 CF in liquid broth and plate mating (Fig. 1A). For R6K and R388, Co4 significantly reduced their CF in both conditions, but the reduction in CF was more pronounced in solid agar mating (R6K, *p* = 0.0062 and R388, *p* = 0.0048) (Fig 1B and 1C). Additionally, Co6 and Co8 had no significant impact on the CF of R388 in liquid broth (Fig. 1C), but significantly reduced its CF in solid agar mating from 4.19×10^-2^ in DMSO control to 9.71×10^-3^ (*p* = 0.0004) and 6.49×10^-3^ (*p* = 0.0003) in Co6 and Co8, respectively (Table S2). None of the cobalt complexes had a significant impact on the CF of pKM101 in both liquid broth and plate mating (Fig. 1D), indicating that IncN plasmids are possibly not targeted by cobalt complexes.

### Impact of cobalt complexes on the transmission of plasmids carrying extended-spectrum β-lactamase and carbapenemase genes

Next, the impact of the four cobalt complexes on the transmission of clinically relevant plasmids carrying ESBL and carbapenemase genes was determined using a previously developed flow cytometry assay to monitor plasmid transmission (30). The recipient *E. coli* and *K. pneumoniae* strains expressed a chromosomal *mcherry* gene whilst the donor *E. coli* strain carrying the IncK type plasmid pCT or the *K. pneumoniae* strain carrying IncFII type plasmid pKpQIL were tagged with constitutively active *gfp*. In this setup, transconjugant bacteria were measured based on their dual fluorescence of GFP and mCherry proteins. None of the four cobalt complexes had a significant effect on the transmission of pCT*gfp* in *E. coli* compared to DMSO control (Fig. 2A), suggesting this IncK plasmid was not the target of cobalt complexes. On the other hand, all four cobalt complexes significantly reduced the transfer of pKpQIL*gfp* in *K. pneumoniae* compared to DMSO control (Fig. 2B).

**Figure 2.**
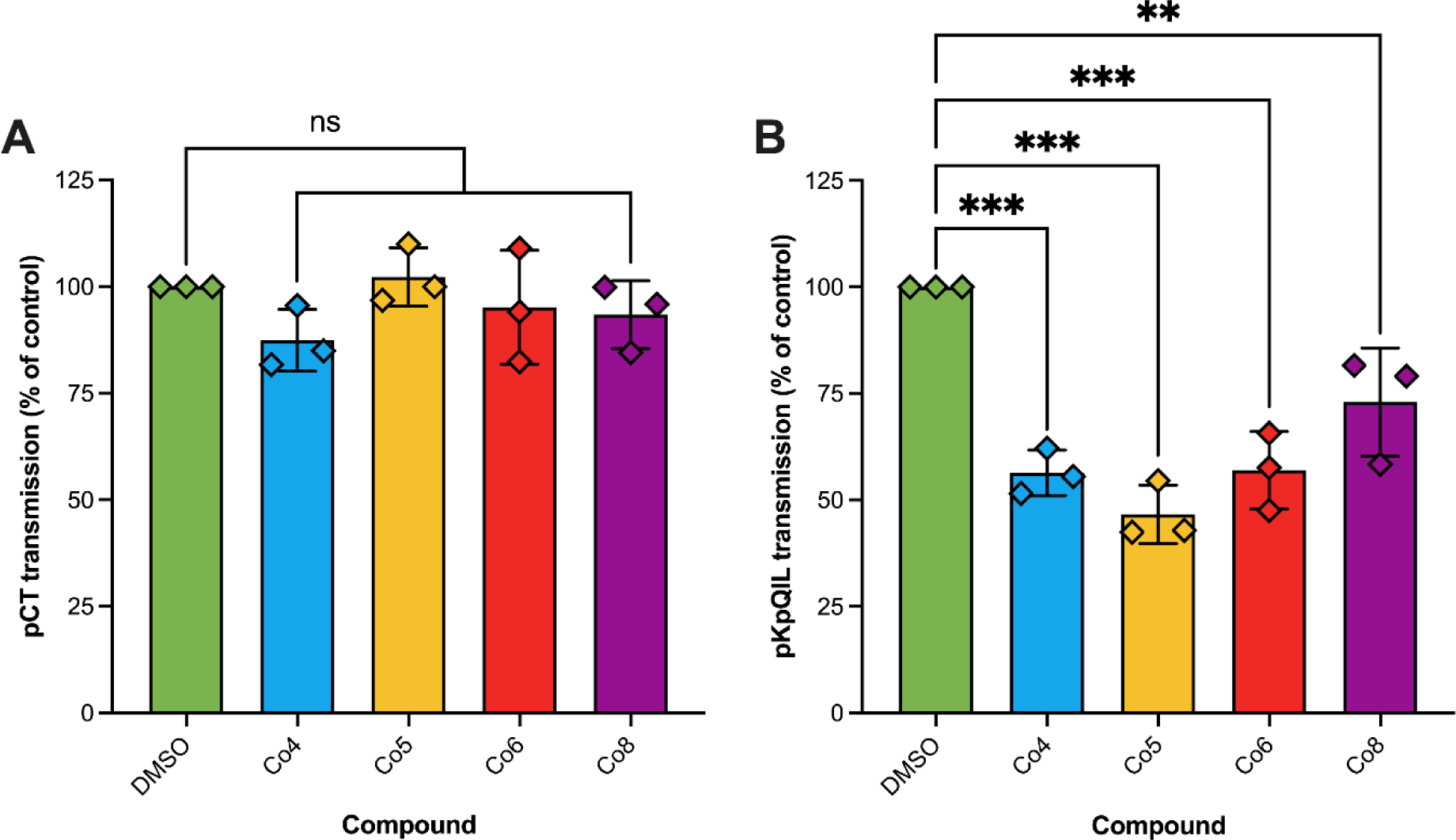
The impact of cobalt complexes on the transmission of plasmids carrying extended spectrum β-lactamase and carbapenemase genes measured by flow cytometry. The transmission of **(A)** pCT*gfp* in *Escherichia coli* ST131 EC958 and **(B)** pKpQIL*gfp* in *Klebsiella pneumoniae* Ecl8, incubated with 100 µg/mL cobalt compound compared to 100 µg/mL DMSO vehicle control after 4 and 6 h incubation, respectively. Data shown are the mean ± standard deviation from three independent experiments, each carried out with four biological replicates. Cobalt complexes that significantly reduced plasmid transmission compared to DMSO control are indicated with ** (*p* ≤ 0.01) or *** (*p* ≤ 0.001). ns, not significant.

### Cobalt complexes do not affect plasmid persistence

To determine whether the cobalt complexes affected plasmid stability and maintenance, the impact of the cobalt complexes on plasmid persistence was measured over 48 h. The DMSO (100 µg/mL) control did not affect plasmid maintenance after 24 and 48 h (Fig. 3). The cobalt complexes did not significantly affect the persistence of all six plasmids compared to DMSO control after 24 and 48 h (Fig. 3). This suggested that the cobalt complexes affected the conjugation process rather than plasmid maintenance in the donor bacteria.

**Figure 3.**
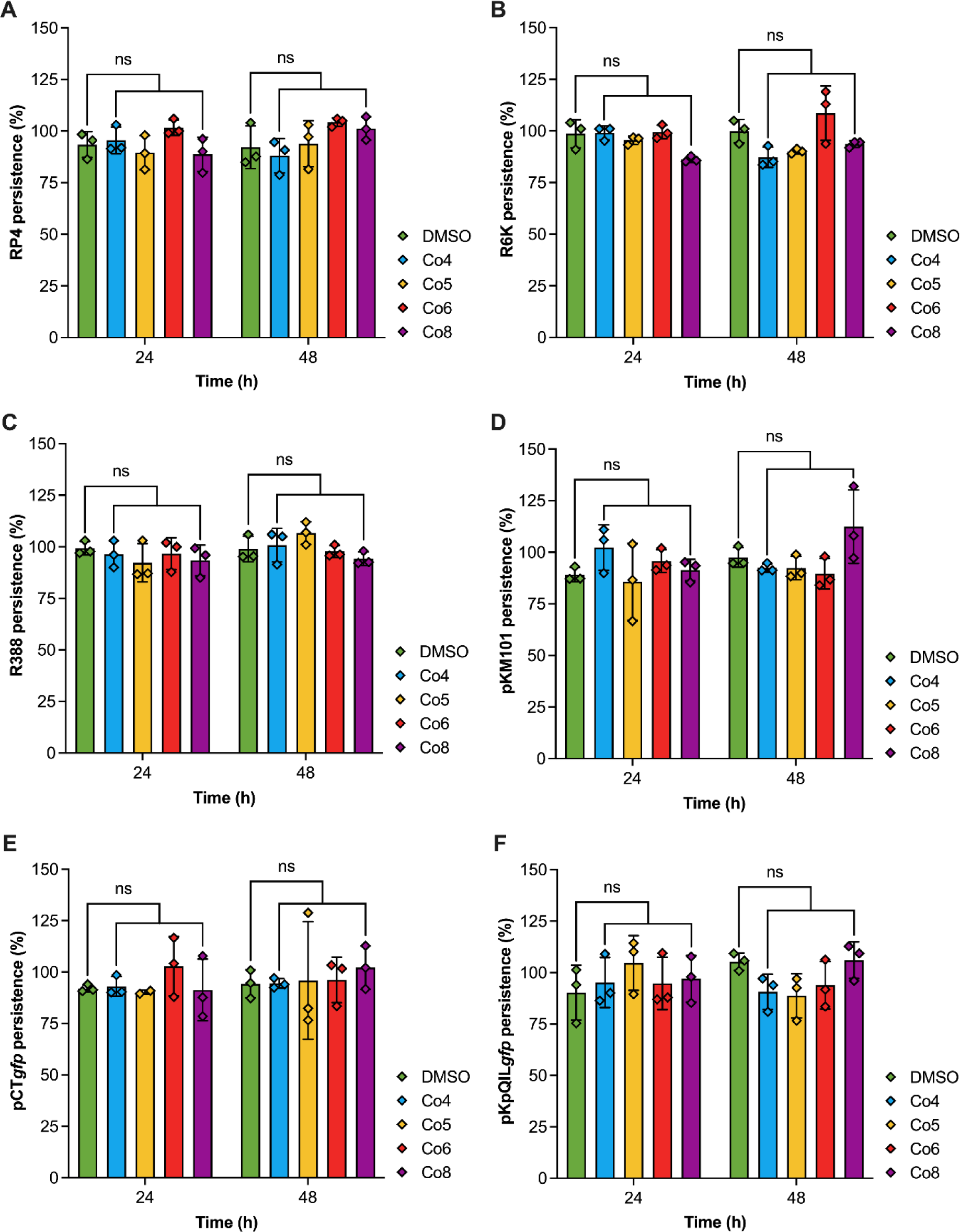
The effect of cobalt complexes on plasmid persistence. The persistence of **(A)** the IncP plasmid RP4, **(B)** the IncX2 plasmid R6K, **(C)** the IncW R388, **(D)** the IncN plasmid pKM101, **(E)** the IncK plasmid pCT with tagged with a *gfp* gene, and **(F)** the IncFII plasmid pKpQIL tagged with a *gfp* gene, in the presence of 100 µg/mL of cobalt complexes after 24 and 48 hours compared to 100 µg/mL DMSO control. Data shown are the mean ± standard deviation from a minimum of three independent experiments, each carried out with four biological replicates. ns, not significant.

## Discussion

The rise in AMR combined with the dwindling pipeline of new antibiotics in development warrants novel strategies to combat the AMR crisis (2). Plasmids play a key role in the global dissemination of AMR genes in MDR Gram-negative bacteria (3). Targeting plasmids is a novel strategy to combat AMR by reducing the prevalence of AMR genes and sensitising bacteria to existing antibiotics (25). In addition, such complexes could be used in a One-Health setting by removing or reducing AMR genes in animals and the environment (3).

Metal ion complexes represent an increasing trend in the development of antimicrobial agents (47). Cobalt complexes have essential biochemical functions and have been reported to possess antibacterial, antifungal, and antiviral properties (37, 48). However, the impact of cobalt complexes on conjugative plasmid transfer has never been explored. In this study, four previously characterised bis(*N*-picolinamido)cobalt(II) complexes (Co4, Co5, Co6, and Co8) were assessed for their ability to reduce conjugation and persistence of different plasmids. The results showed that the complexes did not affect plasmid persistence, suggesting that they affected the conjugation process rather than plasmid stability.

The cobalt complexes were most effective at reducing plasmid conjugation on solid agar, rather than in liquid mating assays. Complexes Co4, Co5, and Co6 significantly reduced RP4 conjugation on solid agar but not in liquid broth (Fig. 1). Similarly, Co4, Co6, and Co8 significantly reduced R388 conjugation on solid agar but only Co4 reduced R388 conjugation in liquid broth as well (Fig. 1). The IncP plasmid RP4 and the IncW plasmid R388 have been previously shown to have constitutive rigid pilus synthesis that plays an important role in conjugation on solid surfaces (45). Cobalt complexes that reduced conjugation specifically on solid agar may target plasmid-specific pilus formation/assembly to impede donor-recipient contact and transfer of single stranded plasmid DNA (49). For the IncX2 plasmid R6K, only Co4 significantly reduced its conjugation frequency in both liquid broth and on solid agar (Fig. 1). Indeed, Co4 reduced solid agar conjugations of 3/4 plasmids, with only pKM101 (which was not affected by any compound in solid or liquid assays) showing no impact. It is possible that Co4 may target a shared component of the conjugation apparatus of these plasmids. It was also the only cobalt complex that reduced R6K conjugation.

All four cobalt complexes significantly reduced the transmission of the IncFII plasmid pKpQIL in *K. pneumoniae* as measured by flow cytometry (Fig. 2B). Therefore, it is plausible that they have a common target that is necessary for successful conjugative plasmid transfer, such as the type 4 secretion system (50), or a common effect on *Klebsiella*. The cobalt complexes do not have any impact on growth kinetics of both *Klebsiella* strains.

The cobalt complexes had no significant effect on the pKM101 conjugation frequency in both solid agar and liquid broth (Fig. 1D), and they also had no effect on the transmission of the IncK plasmid pCT in *E. coli* as measured by flow cytometry (Fig. 2A). These results suggested that the cobalt complexes did not target IncK and IncN plasmids. This is possibly due to diverse elements present in the conjugation apparatus between different plasmid incompatibility groups (44, 51–53).

None of the cobalt complexes exhibited antibacterial activity (Table 3), which corroborates with the previously reported data (37). Moreover, the previous study demonstrated that these cobalt complexes have no cytotoxicity towards mammalian cells (37). To date, this is the first description of cobalt complexes that reduced conjugative plasmid transfer. Further work involving structural modification and mechanism of action studies on bis(*N*-picolinamido)cobalt(II) complexes could potentially lead to the development of broad range conjugation inhibitors.

## Transparency declaration

None to declare.

