## Supplemental Data for "Metal complexes and conjugation: Harnessing the power of cobalt complexes to curtail plasmid transfer"

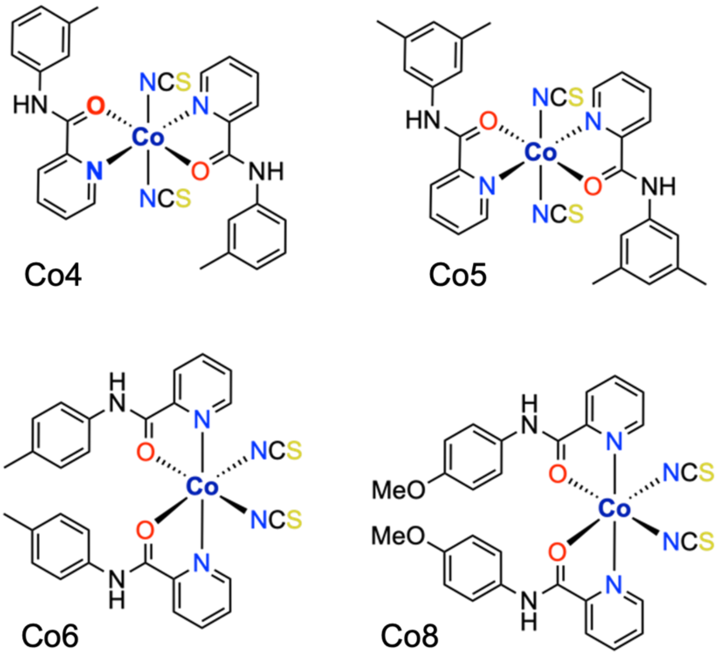


**Figure S1.** The structures of bis(*N*-picolinamido)cobalt(II) complexes Co4, Co5, Co6, and Co8. All complexes have been drawn with the geometries obtained from single crystal X-ray diffraction data (1).

**Table S1.** Primers used in this study.

| Primer ID | Description | Sequence (5'-3') |
|---|---|---|
| P77 | Forward primer for generation of the hygromycin resistance cassette from pSIM18 with flanking 40 bp homology (underlined) to the target locus of the *Tn*7-transposon downstream of the *glmS* gene | TCGGTTACGGTTGAGTAATAAATGGATGCCCTGCGTAAGCGCATTCAAATATGTATCCGCT |
| P78 | Reverse primer for generation of the hygromycin resistance cassette from pSIM18 with flanking 40 bp homology (underlined) to the target locus of the *Tn*7-transposon downstream of the *glmS* gene | AGGATGTTTGATTAAAAACATAACAGGAAGAAAAATGCCCCTATTCCTTTGCCCTCGGACG |
| P87 | Forward primer that binds upstream of the inserted hygromycin resistance gene in the *E. coli* J53 chromosome | GCACCGAACAACGAATTGCT |
| P88 | Reverse primer that binds downstream of the inserted hygromycin resistance gene in the *E. coli* J53 chromosome | TGTTTTACGCCACCGGAAGA |
| P153 | Forward primer that binds to *oriT* on the RP4 plasmid | CAGGATTCCCGTTGAGCACC |
| P154 | Reverse primer that binds to *oriT* on the RP4 plasmid | TCTTTGGCATCGTCTCTCGC |
| P155 | Forward primer that binds to *rep1* on the pKM101 plasmid | GTTCTTCTGTTGGGATTCCG |
| P156 | Reverse primer that binds to *rep1* on the pKM101 plasmid | GCGAAGATGATGATGAGATGG |
| P157 | Forward primer that binds to γ *ori* on the R6K plasmid | CTAAGGGCTTCTCAGTGCGT |
| P158 | Reverse primer that binds to γ *ori* on the R6K plasmid | CGCTATATTACCCCAAACCCG |
| P159 | Forward primer that binds to *repA* on the R388 plasmid | CAGGAACACGCGATAGGTCA |
| P160 | Reverse primer that binds to *repA* on the R388 plasmid | TCACCTTGTCGATCATGGGC |

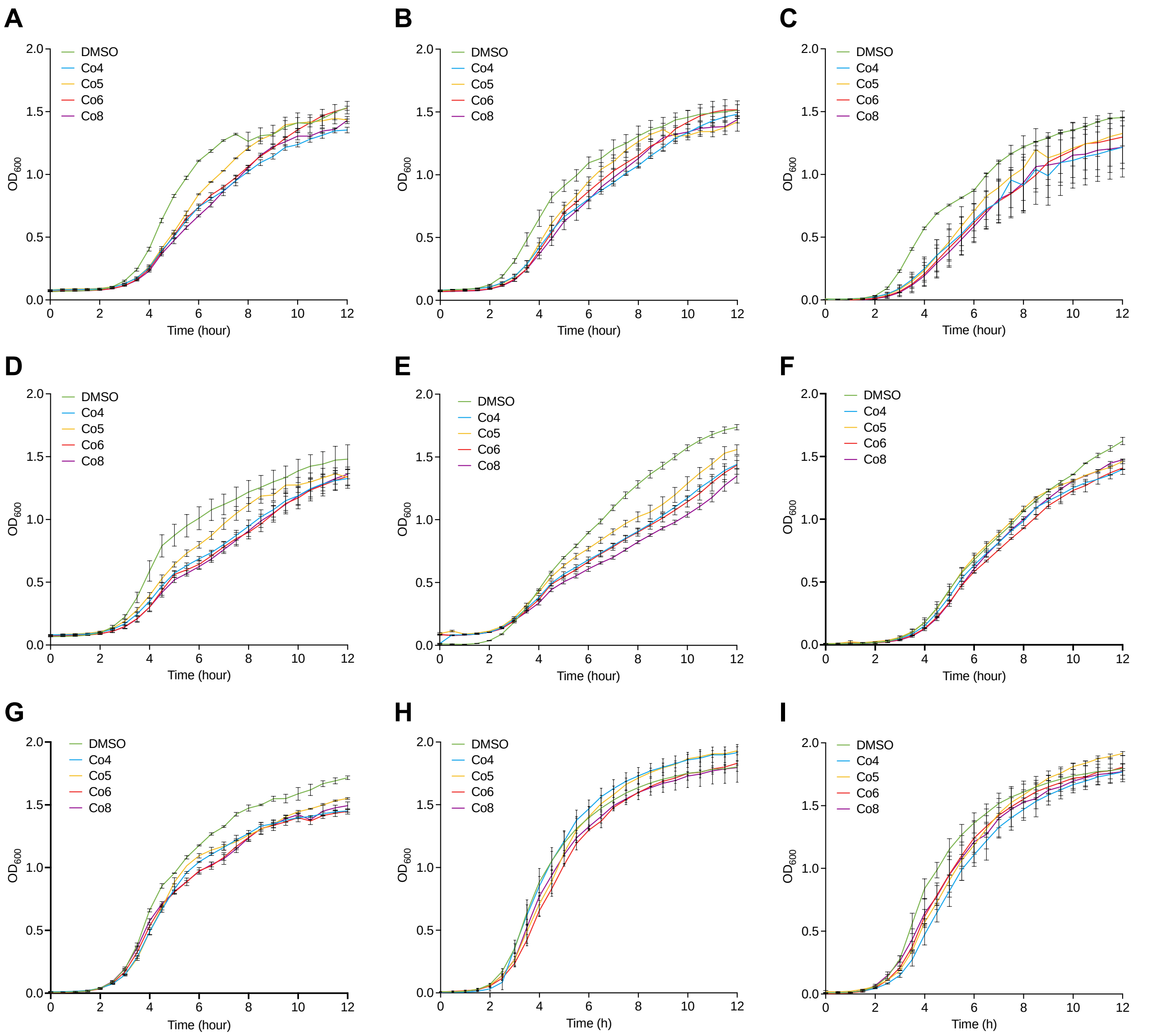


**Figure S2.** The effect of cobalt compounds on growth of bacterial strains. Growth kinetics of **(A)** hygromycin resistant *E. coli* J53 *att*Tn*7*::*hph*, **(B)** *E. coli* J53 carrying IncP plasmid RP4, **(C)** *E. coli* J53 carrying IncX2 plasmid R6K, **(D)** *E. coli* J53 carrying IncW R388, **(E)** and *E. coli* J53 carrying the IncN plasmid pKM101, **(F)** *K. pneumoniae* Ecl8 mCherry, **(G)** *K. pneumoniae* Ecl8 carrying pKpQIL*gfp*, **(H)** *E. coli* ST131 carrying pCT*gfp,* and **(I)** *E. coli* ST131 mCherry, in the presence of LB broth supplemented with 100 µg/mL of cobalt compounds or 100 µg/mL DMSO. Data shown are the mean ± standard deviation of three independent experiments, each carried out with three biological replicates.

**Table S2.** Raw conjugation frequency data for the RP4, R6K, R388 and pKM101 plasmids in plate (solid) and liquid broth mating in the presence of 100 µg/mL of cobalt complexes or DMSO vehicle control.

| | **Conjugation frequency** | | | | | | | |
|---|---|---|---|---|---|---|---|---|
| | **RP4** | | **R6K** | | **R388** | | **pKM101** | |
| Compound | Solid | Liquid | Solid | Liquid | Solid | Liquid | Solid | Liquid |
| **DMSO** | $4.26{\times}10^{-3} \pm 1.1{\times}10^{-3}$ | $4.26{\times}10^{-3} \pm 1.1{\times}10^{-3}$ | $1.19{\times}10^{-3} \pm 4.59{\times}10^{-4}$ | $6.26{\times}10^{-5} \pm 3.62{\times}10^{-5}$ | $4.19{\times}10^{-2} \pm 5.3{\times}10^{-3}$ | $2.11{\times}10^{-3} \pm 1.02{\times}10^{-3}$ | $4.33{\times}10^{-1} \pm 2.93{\times}10^{-1}$ | $1.06{\times}10^{-2} \pm 2.44{\times}10^{-3}$ |
| **Co4** | $6.79{\times}10^{-2} \pm 1.08{\times}10^{-2}$ | $9.06{\times}10^{-3} \pm 7.39{\times}10^{-4}$ | $2.95{\times}10^{-5} \pm 2.56{\times}10^{-5}$ | $1.58{\times}10^{-5} \pm 7.4{\times}10^{-6}$ | $1.82{\times}10^{-2} \pm 6.85{\times}10^{-3}$ | $5.63{\times}10^{-4} \pm 2.66{\times}10^{-4}$ | $3.99{\times}10^{-1} \pm 4.73{\times}10^{-1}$ | $1.08{\times}10^{-2} \pm 1.24{\times}10^{-3}$ |
| **Co5** | $4.5{\times}10^{-2} \pm 3.85{\times}10^{-2}$ | $1.11{\times}10^{-2} \pm 1.57{\times}10^{-3}$ | $6.52{\times}10^{-4} \pm 4.99{\times}10^{-4}$ | $6.53{\times}10^{-5} \pm 2.12{\times}10^{-5}$ | $6.23{\times}10^{-2} \pm 4.02{\times}10^{-2}$ | $1.33{\times}10^{-3} \pm 6.26{\times}10^{-4}$ | $4.97{\times}10^{-1} \pm 3.96{\times}10^{-1}$ | $1.14{\times}10^{-2} \pm 1.54{\times}10^{-3}$ |
| **Co6** | $1.62{\times}10^{-2} \pm 3{,}91{\times}10^{-3}$ | $8.42{\times}10^{-3} \pm 3.51{\times}10^{-3}$ | $7.59{\times}10^{-4} \pm 1.16{\times}10^{-4}$ | $1.63{\times}10^{-4} \pm 3.8{\times}10^{-5}$ | $9.71{\times}10^{-3} \pm 3.2{\times}10^{-3}$ | $1.29{\times}10^{-3} \pm 2.86{\times}10^{-4}$ | $2.81{\times}10^{-1} \pm 1.88{\times}10^{-1}$ | $1.32{\times}10^{-2} \pm 1.56{\times}10^{-3}$ |
| **Co8** | $2.95{\times}10^{-1} \pm 2.84{\times}10^{-1}$ | $7.11{\times}10^{-3} \pm 1.66{\times}10^{-3}$ | $8.78{\times}10^{-4} \pm 4.39{\times}10^{-4}$ | $1.92{\times}10^{-4} \pm 1.01{\times}10^{-4}$ | $6.49{\times}10^{-3} \pm 2.61{\times}10^{-3}$ | $1.2{\times}10^{-3} \pm 3.65{\times}10^{-4}$ | $1.70{\times}10^{-1} \pm 1.01{\times}10^{-1}$ | $1.24{\times}10^{-2} \pm 1.27{\times}10^{-3}$ |

Conjugation frequencies shown are the mean of three independent experiments ± standard deviation, each carried out with four biological replicates.

Conjugation frequency was calculated as number of transconjugants/(number of recipients × donor/recipient ratio).
